# RFP-Cyanine Dye Probe Pair for *In vivo* Neurovascular Multiphoton Imaging

**DOI:** 10.1101/2023.05.04.539515

**Authors:** Fei Xia, David Sinefeld, Zong Chang, Xiaojing Gong, Qinchao Sun

## Abstract

*In vivo* imaging of the neurovascular network is considered to be one of the most powerful approaches for understanding brain functionality. Nevertheless, simultaneously imaging the neuron network and blood vessels in deeper brain layers in a non-invasive manner remains to be a major challenge due to the lack of appropriate labeling fluorescence probe pairs. Herein, we proposed a 2P and 3P fluorescence probe pair for neurovascular imaging. Specifically, the red fluorescence protein (RFP) with an absorption maximum around 550 nm is used as a 3P excited probe to label neurons, and a cyanine derivative dye Q820 has a NIR absorption maximum of 825 nm as a 2P excited probe to label the vasculature, enabling single wavelength excitation at 1650 nm for neurovascular imaging. In particular, the two-photon cross section of Q820 was found to be about 2-fold higher than that of indocyanine green (ICG), a commonly used red two-photon fluorescence labeling agent, at the same excitation wavelength. Benefited from the long wavelength advantage in reducing scattering in both 2 and 3-photon excitation of the fluorescence pairs, we demonstrated in vivo neurovascular imaging in intact mouse brains through white matter and deep into the hippocampus in somatosensory cortex.

## Introduction

Recognized as one of the most intricate and intelligent systems on Earth, the neurovascular network has been at the center of extensive scientific efforts aimed at understanding its complex mechanisms of communication and cooperation. ^1–4^ It has been discovered that there is a profound coordination between neural activity and vascular function. A prominent example of this relationship is neurovascular coupling (NVC), a process whereby capillary blood flow is modulated by neural activity. ^1,2,5,6^ In the quest to elucidate NVC, in vivo imaging techniques have proven indispensable. Functional MRI, which exploits the coupling between neuronal activity and brain hemodynamics, facilitates non-invasive localization and measurement of brain activity.^7^ However, while functional MRI can provide large-scale in vivo brain images, it is constrained by its limited spatial resolution. Conversely, in vivo optical fluorescence microscopy has been developed to visualize biological processes with superior spatial and temporal resolutions. ^8^ Despite these advancements, the technique is challenged when imaging deeper within highly scattered tissues while maintaining high spatial resolution. A variety of innovative imaging techniques, primarily employing long-wavelength excitation to diminish tissue scattering, have been proposed to overcome this limitation.^9–11^ Among these innovations is the short-wave infrared (SWIR) one-photon fluorescence imaging, developed to leverage the advantages of long emission wavelengths for deep *in vivo* tissue imaging. ^12^ Moreover, long wavelength/SWIR two-photon (2P) and three-photon (3P) fluorescence imaging have been utilized for *in vivo* 3D deep mouse brain imaging via long-wavelength excitation. ^13,14^

For long wavelength fluorescence microscopy, reliable and bright fluorescent probes are essential for intravital deep imaging for biological studies.^15^ For instance, the toolbox development of fluorescence protein makes the *in vivo* labeling neuron network feasible.^16^ Thanks to the significant 3P absorption cross section of fluorescence proteins, *in vivo* 3P imaging of the fluorescence protein labeled neuron network has been demonstrated reaching over 1 mm in deep mouse brain.^13,14^ However, in comparison to fluorescent protein, small organic fluorescent molecules possess unique advantages, including easy modification of their photophysical properties and molecular structures. Specifically, the cyanine dyes are one of the most studied fluorescent probes for imaging. For example, ICG is the only FDA-approved dye with emission spectrum greater than 800 nm.^15^ Furthermore, ICG has a reasonable two-photon cross section for *in vivo* imaging.^17^ However, due to its limited brightness and relatively short one-photon absorption peak wavelength, ICG has limited application in deeper brain imaging.

It was demonstrated that the optimal imaging windows for mouse brain imaging are around 1300 nm and 1700 nm, where 1700 nm being most promising for deeper structural imaging.^18^ 3P fluorescence imaging of RFP labeled neuron network has been reported for deep brain tissue with 1700 nm excitation.^13^ However, simultaneously imaging the neuron network and brain vessels in deeper brain layers is still considered to be a great challenge for the lack of right labeling fluorescence probe pair. In previous work, it was demonstrated that neuronal and vascular excitation can be imaged simultaneously at wavelengths at 1300 nm, but one fluorophore must be excited to higher energy levels.^19^ Herein, we aim to propose a generalized strategy to allow multicolor imaging around 1700 nm combining a 2P and 3P fluorescence probe pair, for which the RFP with absorption maximum around 550 nm as the 3P excited probe, and the cyanine derivative dye Q820 with absorption maximum around 825 nm as the 2P excited probe. The long excitation wavelength was chosen at 1650 nm. The two-photon cross section of Q820 was found to be about 2-fold higher than that of ICG at the excitation wavelength. A through white matter brain vessel image was achieved as deep as 1.7 mm via the dye Q820. The neurovascular network was recorded via such proposed fluorescent probe pair.

## Results

### Characterization of RFP, ICG and Q820

The absorption peak of RFP (tdimer2 (12)) is around 550 nm, and the proper 3P excitation wavelength was chosen around 1650 nm, Figure 1b. To perform 2-photon fluorescence imaging with the same excitation wavelength with efficient 2P excitation, a Q820 dye was prepared, whose absorption peak was approximately 825 nm. To obtain a well-biocompatible fluorescent probe, the Q820 was incubated first with BSA to form a Q820@BSA complex, Figure 1a. The absorption and emission spectra of Q820@BSA, ICG and RFP (tdimer2 (12)) were shown in Figure 1c and 1d. The maximum absorption of Q820@BSA is around 825 nm which is about 40 nm redshifted from that of ICG (785 nm). For the RFP (tdimer2 (12)), the maximum absorption is around 550 nm. We chose an excitation wavelength around 1650 nm for efficient 2P excitation of Q820@BSA and the 3P excitation of RFP. The two and three photon fluorescence as a function of the excitation laser intensity with excitation at 1650 nm was shown in Figure 1e. A slope around 2 for both Q820@BSA and ICG fluorescent and about 3 for RFP was observed. The normalized two-photon fluorescence of ICG and Q820 was shown in Figure 1f. For ICG and Q820@BSA at the same molecular concentration (1 *μ*M), the corresponding two-photon fluorescence intensity was found to be about 2 folds higher for Q820@BSA than ICG at excitation of 1650 nm.

**Figure 1.**
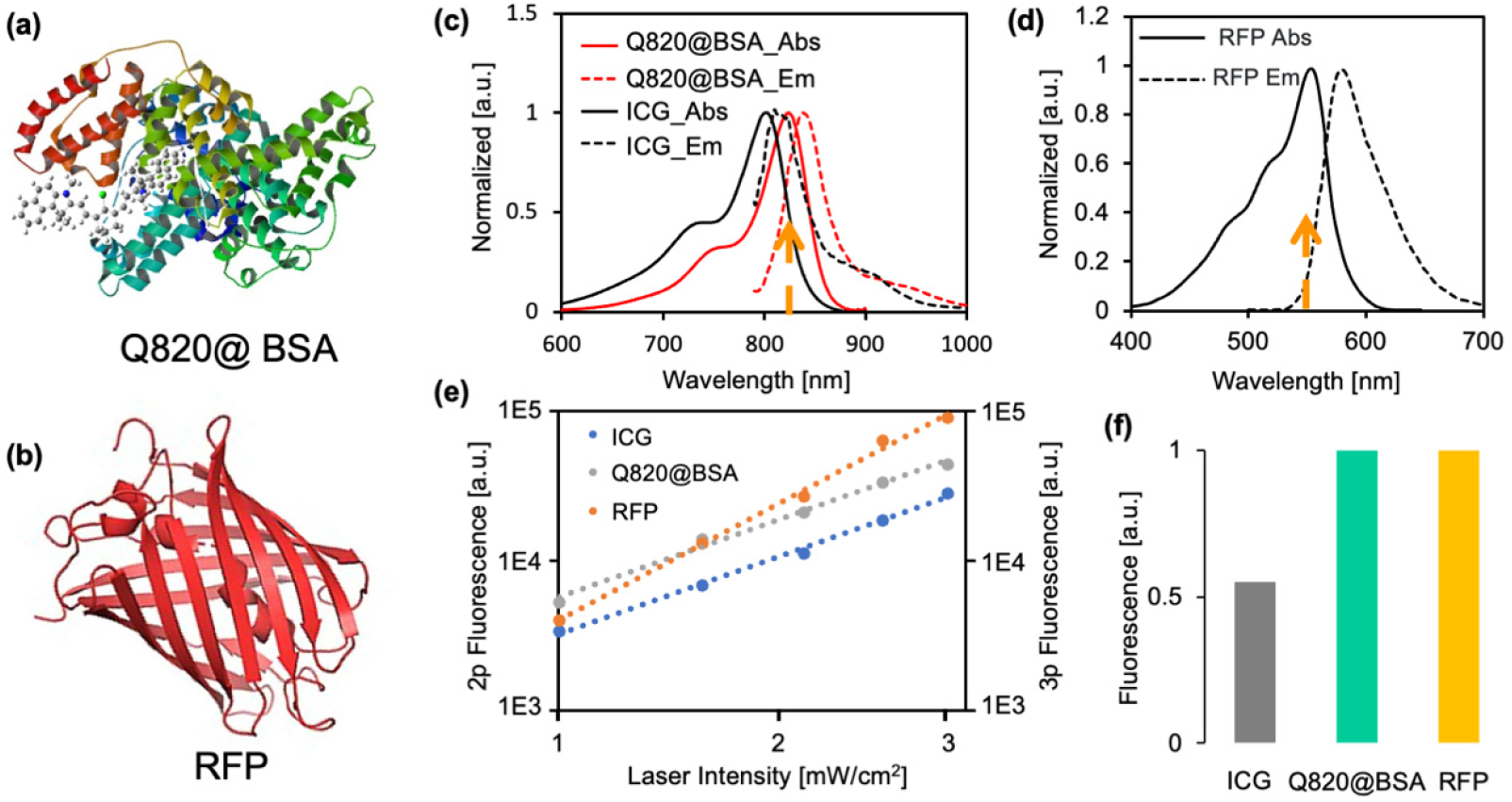
(a) The molecular geometry of Q820@BSA. (b) The molecular geometry of RFP. (c) Absorption and emission spectrum of ICG, Q820@BSA. (d) Absorption and emission spectrum of RFP. (e) (left axis) The two-photon fluorescence intensity as a function of excitation laser intensity of ICG and Q820@BSA; (right axis) the three-photon fluorescence intensity as a function of excitation laser intensity of RFP. All axis were in log scale. (f) The normalized multiphoton fluorescence of ICG and Q820@BSA (2p), RFP (3p) with excitation at 1650 nm.

### Deep *in vivo* 2P fluorescence imaging of vasculature in mouse brain

The 2P fluorescence imaging of the brain vascular network was performed via retro-orbital injection of Q820@BSA and with fluorescence excitation about 1650 nm. A 3D image stack was reconstructed for the 2P fluorescence channel as shown in Figure 2. The THG channel was imaged simultaneously. Providing label free information of the brain tissue. During the first few hundreds of microns deep in the cortical layers, the THG contrast is mainly due to lipid-rich structures such as blood vessels and myelinated axons. Through THG, the white matter could be clear distinguished with the characterized pattern of the filaments from 875 *μ*m to 950 *μ*m as shown in Figure 2 and video 2. THG signal is degraded when imaging deeper. After the whiter matter layer is reached, there is a significant drop in THG signal, and a stronger background is visible in the 2P fluorescence channel as a result of the increased scattering from white matter layers. Despite strong scattering, we were still able to visualize blood vessels at the depth around 1700 *μ*m.

**Figure 2.**
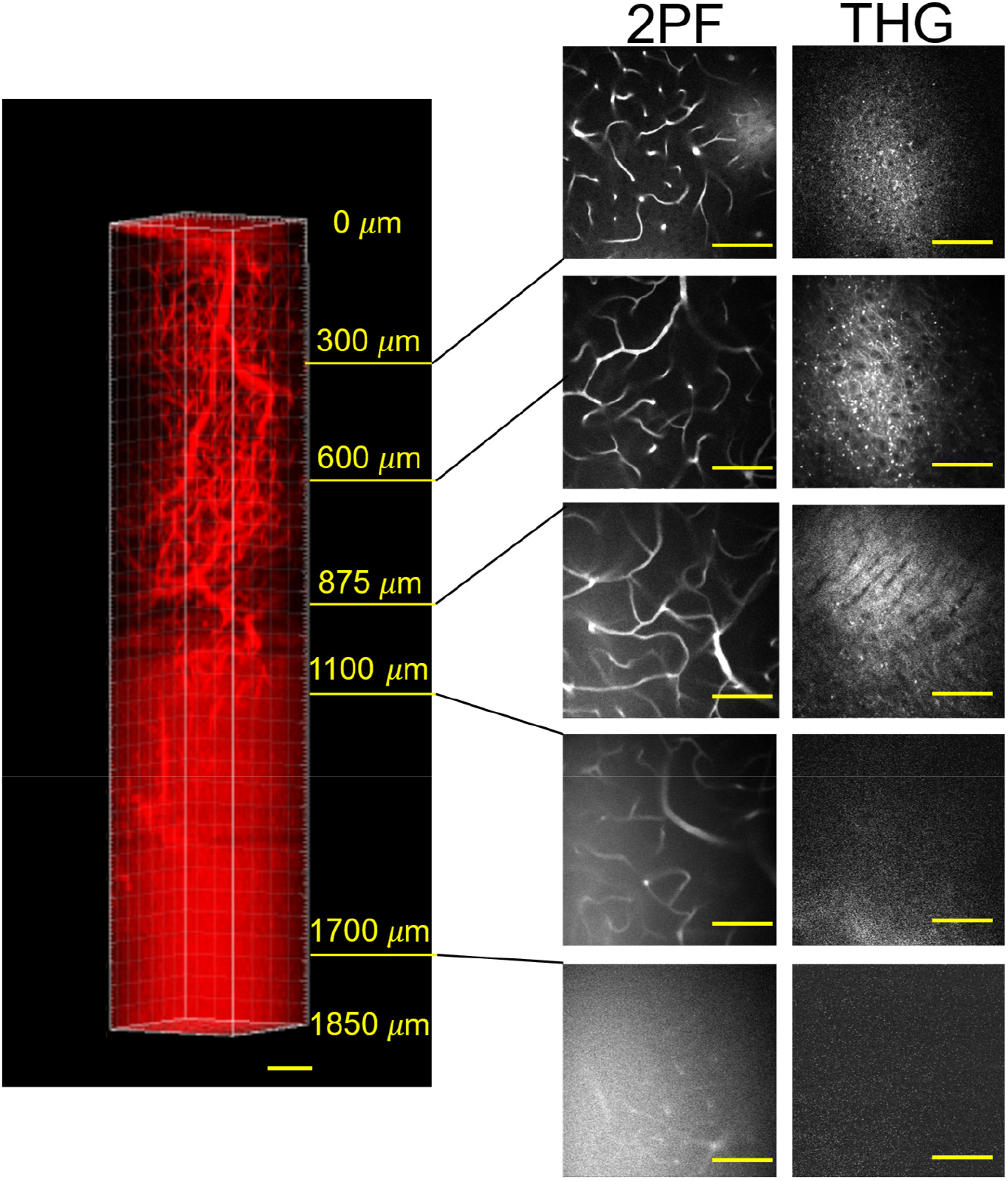
The 3D reconstruction of the brain vascular network via 2-photon fluorescence imaging with excitation at 1650 nm, and the corresponding third-harmonic generation (THG) images with scale bar at 100 *μ*m.

### Characterization of 2P fluorescence imaging with Q820

The effectiveness attenuation length was used to characterize the optical properties of the brain tissue in order to quantitatively evaluate how deep we had penetrated in the mouse brain. The attenuation THG signal as a function of image depth was analysed, instead of the two-photon fluorescence channel. In comparison to the intrinsic probe of THG, the two-photon fluorescence signal could be affected by the probe properties, for instance the blood circulation time. Each image’s signal was calculated as the average value of the top 0.1% of all pixels. The natural logarithm of THG signal normalized by cubic square of power used at the brain surface *S*/*P*^3^ as a function of the imaging depth was shown in Figure 3. In a heavily scattering tissue environment, the intensity of the excitation beam at the focus would be greatly reduced as a function of the imaging depth. Since the fluorescence signal was collected by a large area bucket detector (i.e. PMT), the scattering loss of fluorescence signal could be negligible. The recorded multiphoton fluorescence signal would be mostly dominated by the signal attenuation of the excitation beam. The ln 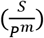 as a function of the imaging depth could be described as Equation 1, where *P* is the power used at the brain surface for each imaging depth, m is the order of nonlinear process, *C* is a constant and *l_e_* is effective attenuation length.

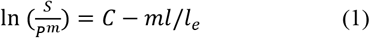

A clear turning point could be observed at the depth around the 900 *μ*m in Figure 3. According to equation 1, the change in the slope corresponds to the variation of the effective attenuation length *l_e_*. It is consistent with the adult mouse brain anatomy that the low scattering somatosensory cortex has a thickness of about 1 mm, and beyond that the high scattering white matter is present. Under the white matter lies the hippocampus region. The *l_e_* of cortex was found to be around 370 *μ*m. The white matter exhibits much smaller *l_e_* as about 179 *μ*m. There is no observable THG signal from the hippocampus tissue.

**Figure 3.**
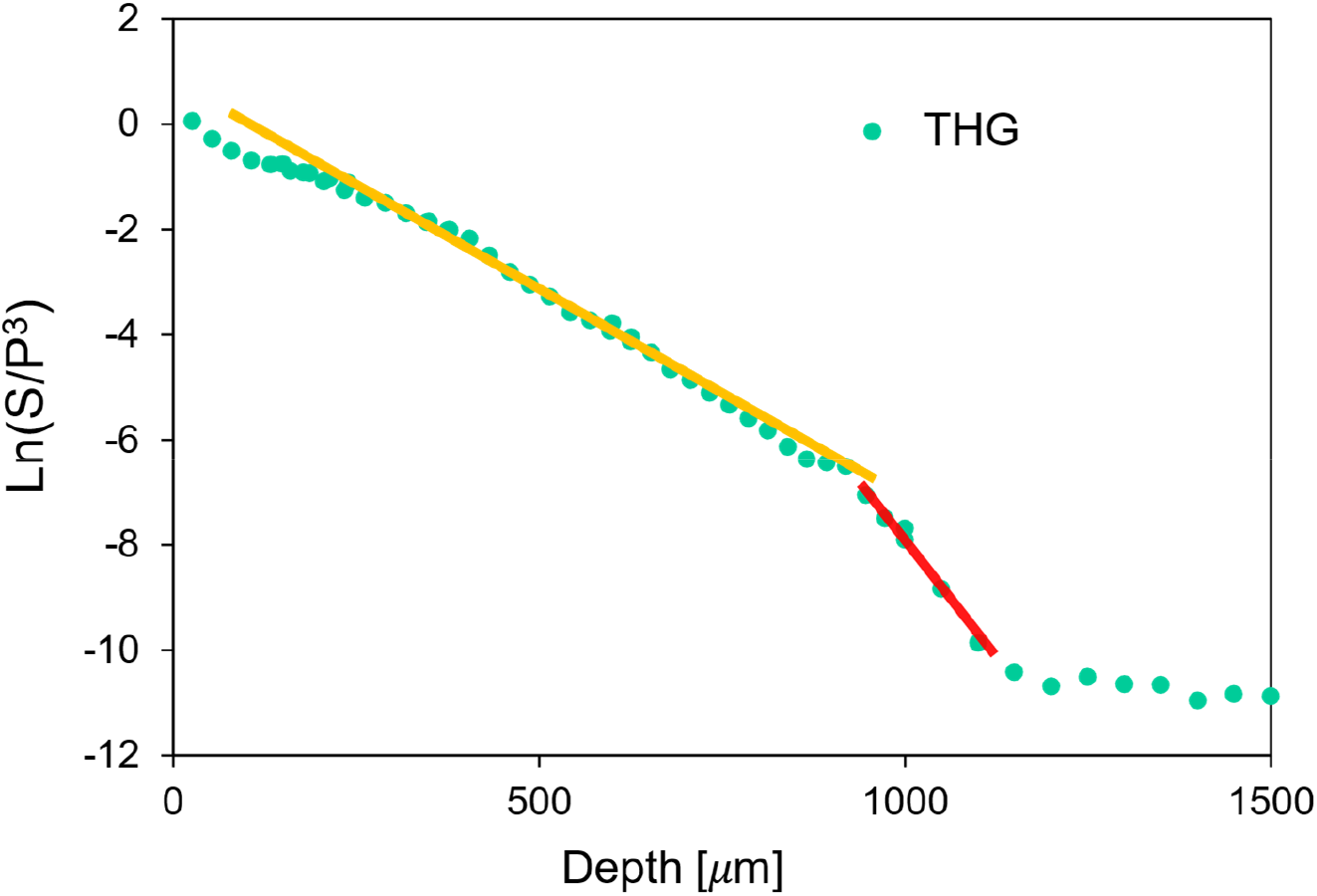
THG signal decay curve plotted on a semi-log scale for calculating the effective attenuation lengths in the neocortex and the white matter layer. The straight lines are linear fitting results for calculating the effective attenuation length *l_e_*.

The signal to noise ratio (S/N) of the 2P fluorescence image at depth around 1000 *μ*m within white matter and 1600 *μ*m within hippocampus is shown in Figure 4. The noise is calculated by the standard deviation of the background signal and the signal is subtracted from the background. The S/N ratio at the image depth around 1000 *μ*m was about 20, and about 7 at around 1600 *μ*m. The signal-to-background ratio (SBR) is around 6.5 at 1600 *μ*m, which is far beyond the depth limit of fluorescence microscopy (SBR=1). It is also clearly shown in the images that high-resolution capillary is still visible with 2P vasculature imaging with Q820 at 1650 nm excitation.

**Figure 4.**
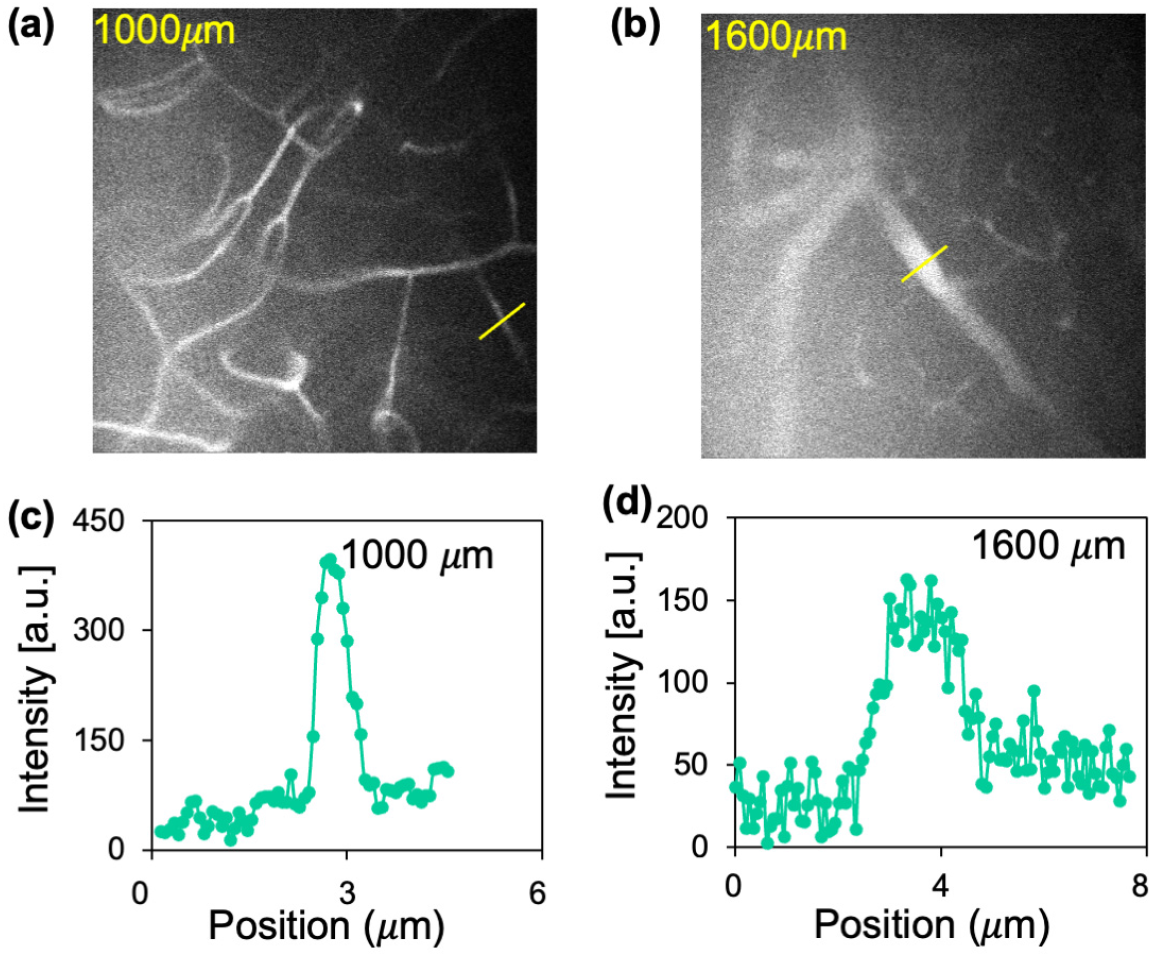
The signal to noise ratio of 2P fluorescence image. (a) 2P fluorescence image of brain vessel at depth about 1000 *μ*m. (b) 2P fluorescence image of brain vessel at depth about 1600 *μ*m. The upper left labeled numbers in (a,b) indicate the depth in the unit of microns. (c) The 2P fluorescence profile of the region of interest in (a) yellow line. (d) The 2P fluorescence profile of the region of interest in (b) yellow line.

### Simultaneous deep neuron and vasculature imaging within living intact mouse brain

Although previous work has shown simultaneous multi-color imaging of neurons and vasculature with 3-photon excitation, it either requires complicated custom laser sources or a specific excitation scheme.^19,21^ For deeper imaging, longer excitation wavelengths of 1700 nm are preferred. It should be noted, however, that *in vivo* imaging in the 1700 nm window is limited by the availability of bright and biocompatible fluorophores. As part of our research, we developed and applied Q820 probes for labeling mouse vasculature. Using excitation around 1650 nm, deep 2-photon imaging was possible through white matter, as demonstrated above. Moreover, the 3P photon excitation of RFP occurs within the same excitation region. We have also proposed and achieved neurovascular imaging using Q820-RFP probe pairs excited at 1650 nm shown in Figure 5, and the corresponding video as supplementary video 1. This is achieved by simultaneous 2-photon excitation of Q820 for vasculature imaging and 3-photon excitation of RFP for neuronal imaging, at the same wavelength around 1650 nm in the 1700 nm imaging window. The neuron cells could be observed up to 1200 *μ*m at depth, limited by the brightness of RFP, while clear vessels could be observed much deeper up to 1500 *μ*m through highly scattered white matter layers. Consequently, we anticipate further efforts to be made to develop brighter red fluorescent proteins to enable the imaging of the neurovascular system at greater depths.

**Figure 5.**
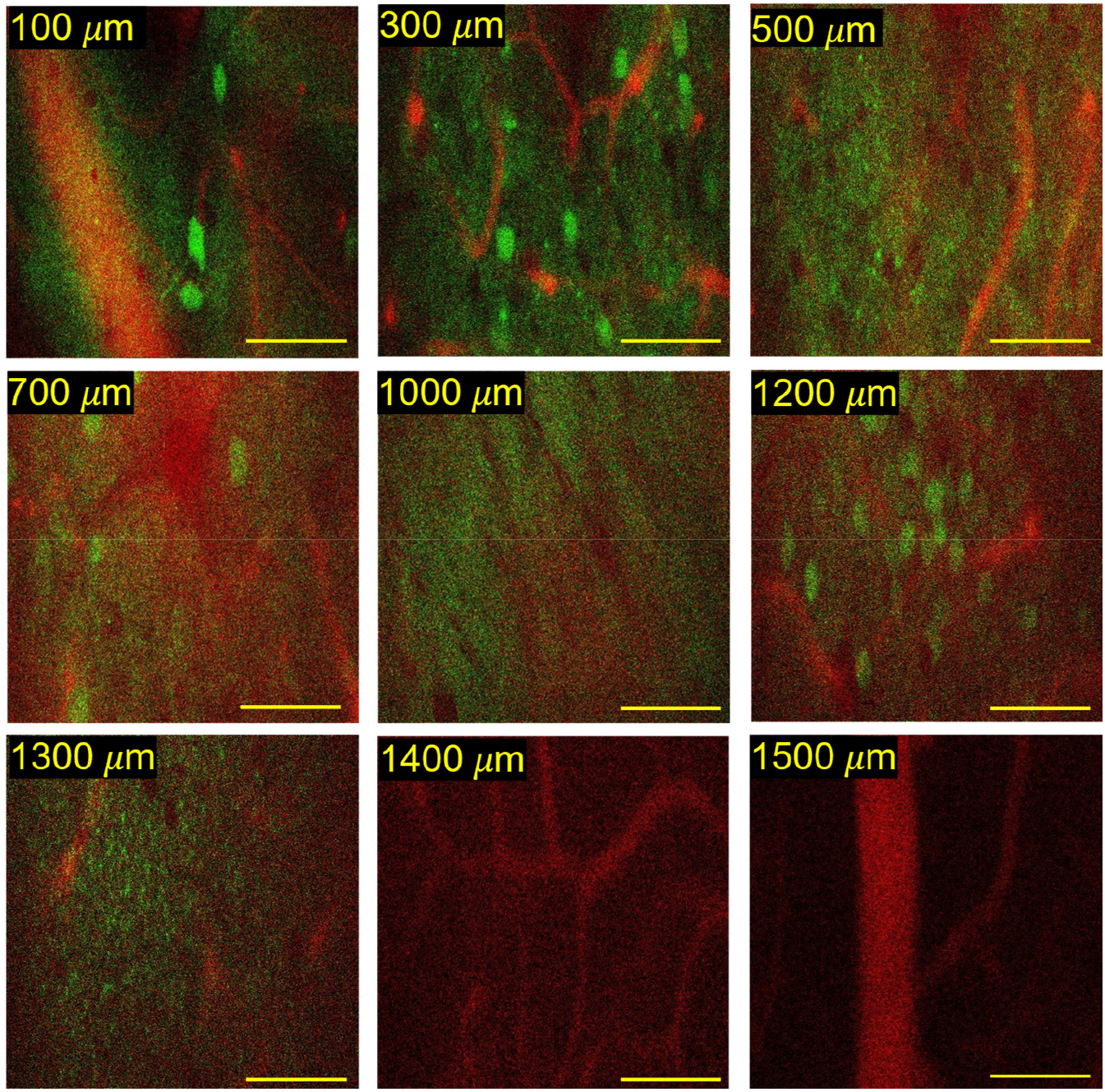
The neurovascular imaging of *in vivo* mouse brain at 1650 nm excitation with 2P fluorescence for Q820 labeled blood vessels and 3P fluorescence for RFP labeled the neurons. Scalar bar: 100 *μ*m.

## Conclusion

The ability to image the neurovascular network in the intact mouse brain holds considerable scientific interest due to its importance in supporting and regulating neuronal activity. Deep imaging of these structures is pivotal as they play critical roles, including nutrient transport, waste removal, and overall neuronal health maintenance. Non-invasive, high-resolution imaging of these deep-lying structures in intact mouse brains can yield invaluable insights into the physiological and pathological processes related to neurovascular coupling. In this study, we have introduced a neurovascular probe pair, RFP and Q820, by matching the transition energy between the three-photon excitation of RFP and the two-photon excitation of Q820, we are able to conduct *in vivo* simultaneous 2- and 3-photon fluorescence structural imaging of neurovascular through the highly scattering white matter in an adult mouse brain excited at the same wavelength. Our investigations also suggest a potentially high two-photon absorption cross-section for Q820, enabling the capture of deep vascular labelling images extending to a depth of 1700 *μ*m beyond the white matter.

Our findings underscore the potential for simultaneous imaging of neuronal cells and brain vessels using three-photon and two-photon mechanisms at the same excitation wavelength, a method that could be broadly applicable due to its inherent generalizability. This method not only streamlines the imaging process but also minimizes potential tissue damage from multiple excitation wavelengths. We hope it provides methodological foundation for future *in vivo* studies investigating neurovascular interactions in deeper brain tissues and stimulates further development of advanced fluorescent probes that are bright, biocompatible, and capable of emitting near-infrared (NIR) fluorescence. The advancement of these tools can improve our capacity to explore the complex interactions within the brain’s neurovascular network, thereby deepening our understanding of its function and contributing to potential strategies for addressing neurological disorders.

## Methods

### Materials

All reagents and solvents were used as received without further purification. Bovine serum albumin (BSA) (98%) was purchased from Sigma-Aldrich (St. Louis, USA). ICG and all other chemicals were bought from Aladdin (Shanghai, China). The extracted RFP was a gift from a biosynthetic lab.

### Preparation of Q820

The synthesis of Q820 was based on the typical cyanine dye preparation procedure. Briefly, compound A was first synthesized by mixing 1,1,2-trimethyl-1H-benzo[e]indole (1.05 g, 5.0 mmol) in toluene (10 mL), EtI (0.48 mL, 6.0 mmol) under argon atmosphere. The reaction mixture was stirred under reflux overnight, then allowed to cool to room temperature. The solvent was removed under vacuum filtration and the residue was washed with ether to afford the desired product 12 (1.5 g, 82%) as a blue solid. To a solution of compound A (182.5 mg, 0.5 mmol) in EtOH (10 mL), sodium acetate (41 mg, 0.5 mmol) and N-((E)-(2-chloro-3-((E)-(phenylimino)methyl)cyclohex-2-en-1-ylidene)methyl)aniline hydrochloride (90 mg, 0.25 mmol) was added gradually under argon atmosphere. The reaction mixture was stirred under reflux for 4 h, then allowed to cool to room temperature. After evaporation of solvent, the residue was purified by flash chromatography. HNMR and HRMS were performed to characterize the final product.

### Imaging setup

We used a wavelength-tunable optical parametric amplifier (OPA, Opera-F, Coherent) pumped by an amplifier (Monaco, Coherent). The laser was operated at 1650 nm. The repetition rate and the optical power were tuned to maintain a similar signal-to-noise ratio at various depths. A laser scanning microscope (Movable Objective Microscope, Sutter Instrument) was used. A silicon wafer was inserted in the beam path to compensate the dispersion in the system.^20^ For the detection of the Q820 fluorescence signal, a GaAs photomultiplier tube (H7421-50, Hamamatsu) and a long pass filter (800 nm) were used. For the detection of the third-harmonic signal, a GaAsP photomultiplier tube (H7421-40, Hamamatsu Photonics) and a 558 ± 10 nm bandpass filter (Semrock) were used. Current generated by the PMT was converted to voltage (0.1 VμA^-1^) and low-pass-filtered (20kHz) by a transimpedance amplifier (C7319, Hamamatsu Photonics). Analog-to-digital conversion was performed by a data acquisition card (NI PCI-6110, National Instruments. We use a custom high NA water immersion microscope objective (XLPLN25XWMP2, Olympus, 25× 1.05 NA) with coating for high transmission (∼80%) at 1,650 nm. A dichroic mirror (750DCXXR, Chroma Technology) was used to direct the fluorescence and third harmonic signal to the PMTs.

### Animal preparation and *in vivo* imaging

Animal procedures were reviewed and approved by the Cornell Institutional Animal Care and Use Committee. We used male wildtype mouse (C57BL/6J, 8 weeks old, Jackson Laboratory) for Q820 labelled vasculature imaging, and B6.Cg-Tg(Thy1-Brainbow1.0) HLich/J mice (18 g, 8 weeks old, The Jackson Laboratory) for RFP-labelled neuron imaging. The B6.Cg-Tg(Thy1-Brainbow1.0) HLich/J mice were not subjected to Cre recombinase, so the only fluorescent protein expression was from RFP tdimer2(12). Craniotomy was performed on an 8-week-old mouse centered at 2 mm posterior and 2 mm lateral to the Bregma point. The cranial window with a circular shaped glass of 5 mm in diameter was mounted directly on the intact dura during imaging. The mouse was immediately imaged after craniotomy and was anesthetized with isoflurane (1-2% in oxygen for induction and 1.5-2% for surgery and imaging to maintain a breathing rate of 1 Hz). During the in vivo imaging session, the mouse body temperature was maintained at 37.5 °C with a temperature-controlled blanket (Harvard Apparatus) and the eyes were kept lubricated with eye ointment. Heavy water (D_2_O) was used as the immersion medium to minimize the absorption of the excitation wavelength at 1650 nm.

For imaging of the vasculature in the mouse brain in vivo, the dye was retro-orbitally injected for labeling. All images were taken at a frame rate of 0.24 Hz (512 × 512 pixels/frame) with a field of view (FOV) of 340 × 340 µm. At the largest depth, the frame was averaged 25 times. For imaging depth measurement, because the reported imaging depths are based on the raw axial movement of the objective, the slightly larger index of refraction in brain tissue relative to water resulted in a slight underestimate (5–10%) of the actual imaging depth within the tissue.^13^

## Acknowledgments

This work was supported by the National Natural Science Foundation of China (no. 22077135) and Natural Science Foundation of Shenzhen (no. JCYJ20200109115633343). We are grateful to Dr. Chris Xu for providing the imaging setup.

## Notes

### Competing Interest Statement

The authors have declared no competing interest.

### Summary of Updates

corrected typos

